# ARBEL: A Machine Learning Tool with Light-Based Image Analysis for Automatic Classification of 3D Pain Behaviors

**DOI:** 10.1101/2024.12.01.625907

**Authors:** Omer Barkai, Biyao Zhang, Bruna Lenfers Turnes, Maryam Arab, David A Yarmolinsky, Zihe Zhang, Lee B Barrett, Clifford J Woolf

## Abstract

A detailed analysis of pain-related behaviors in rodents is essential for exploring both the mechanisms of pain and evaluating analgesic efficacy. With the advancement of pose-estimation tools, automatic single-camera video animal behavior pipelines are growing and integrating rapidly into quantitative behavioral research. However, current existing algorithms do not consider an animal’s body-part contact intensity with- and distance from- the surface, a critical nuance for measuring certain pain-related responses like paw withdrawals (‘flinching’) with high accuracy and interpretability. Quantifying these bouts demands a high degree of attention to body part movement and currently relies on laborious and subjective human visual assessment. Here, we introduce a supervised machine learning algorithm, ARBEL: Automated Recognition of Behavior Enhanced with Light, that utilizes a combination of pose estimation together with a novel light-based analysis of body part pressure and distance from the surface, to automatically score pain-related behaviors in freely moving mice in three dimensions. We show the utility and accuracy of this algorithm for capturing a range of pain-related behavioral bouts using a bottom-up animal behavior platform, and its application for robust drug-screening. It allows for rapid objective pain behavior scoring over extended periods with high precision. This open-source algorithm is adaptable for detecting diverse behaviors across species and experimental platforms.

## Introduction

Pain is an individualized sensation generated by sensory-discriminative and affective neural pathways. Translational models in rodents that simulate pain through the elicitation of defined behaviors play a vital role in the study of pain mechanisms and advancing analgesic development ^1–3^. A fundamental component of these approaches involves defining, and then accurately capturing, a relevant range of behavioral reactions in these models ^4^. Pain evaluation in rodents traditionally requires a visual assessment and precise timing of distinct behavioral occurrences, encompassing actions like flinching, licking, and biting. These behaviors entail more detail than behavioral assays based predominantly on an animal’s location in space (e.g., open field test, place conditioning) and therefore require labor-intensive observations, which are slow and susceptible to human error/bias ^5^. Even when conducted by skilled investigators, preclinical pain-behavior scoring is a bottleneck in the experimental pipeline execution. Importantly, subjective assessment of limited behaviors introduces reproducibility challenges, a major obstacle. In consequence, enhancing the objectivity and throughput of the assessment of pain-related behaviors in preclinical models is of paramount significance for both studying pain and discovering novel interventions.

While for certain behaviors, such as paw licking/biting or scratching, a two-dimensional analysis of pose changes in the horizontal plain (*x-y* surface, Figure 1A) from a bottom-up view suffices, other behaviors require the consideration of a third vertical depth dimension (z axis, Figure 1A). A major behavior that involves such consideration is hindpaw flinching, a repetitive paw twitching and elevating reaction in response to the induction of acute nociceptive pain or chronic pain models. In many pre-clinical pain models non-evoked flinching is the most informative behavior measure of ongoing/spontaneous pain and a sensitivity to touch, however, its detection is challenging due to its fast and short occurrence, especially in persistent pain models requiring long duration monitoring ^6–8^.

**Figure 1.**
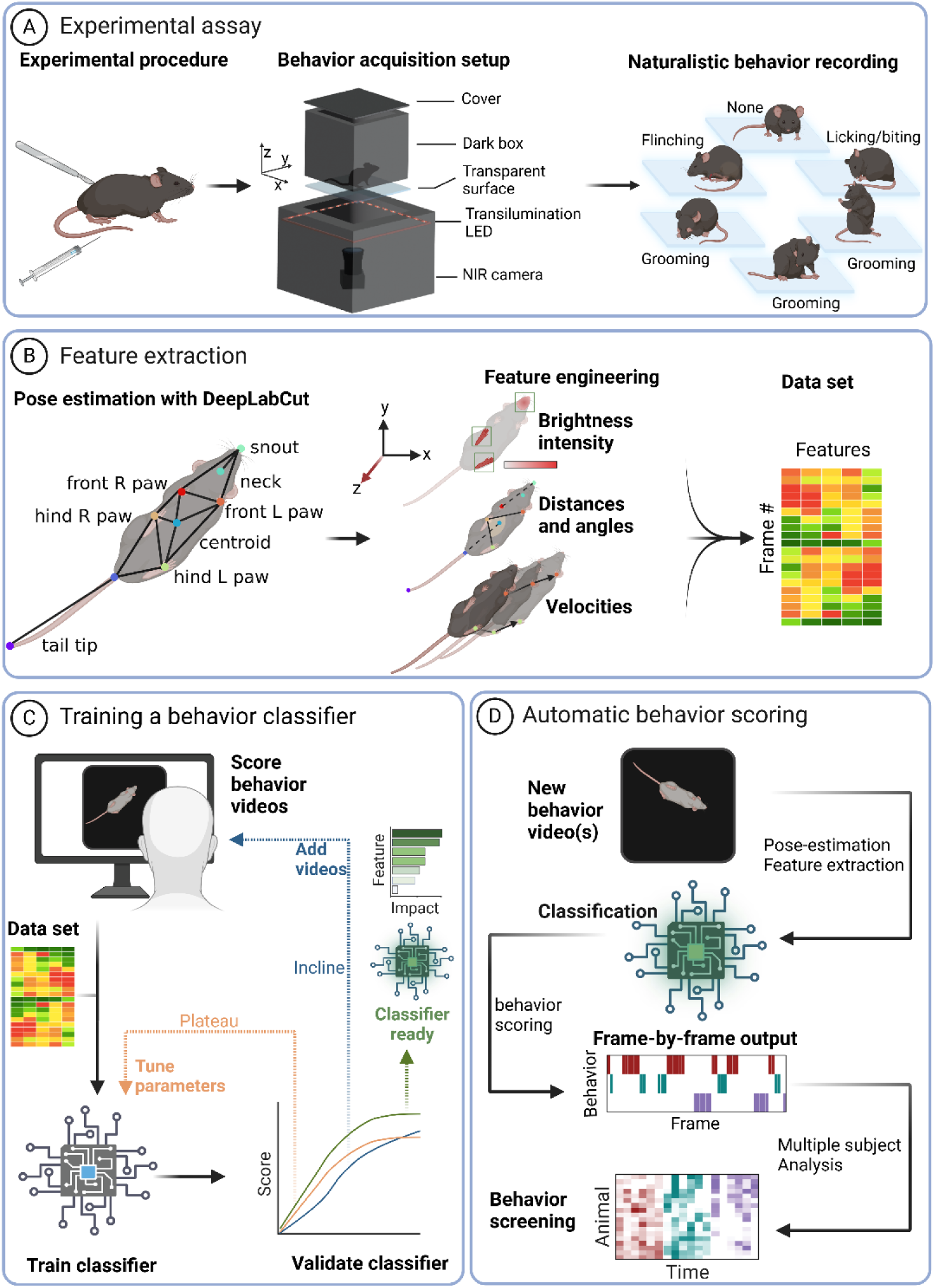
ARBEL workflow from experiment to automated behavior assessment. (A) Diagram of the experimental recording setup. Test mice are put into a behavior platform allowing them to freely move in a dark box on a transparent surface, while a NIR camera together with NIR (∼850nm) LED illumination, captures body movements. The coordinate system defines the surface as the *x-y* plain, and the vertical depth, perpendicular to the surface, along the *z* axis. (B) Illustration of nine key animal body parts labeled for tracking by DLC pose estimation before feature extraction (left). After DLC tracking, three feature types are constructed by frame-by-frame body part coordinates together with video image analysis (middle). Hindpaw and snout brightness capture and quantification for the brightness feature set are illustrated (red color scale). The green rectangles depict regions of interest in which pixel brightness is averaged for quantifying paw contact with the surface. Dashed black lines show example distances, the orange surface between solid black lines illustrates an example angle, measured from pose-estimated body parts. Velocity is calculated as the distance a body part moves over a specified number of frames. The data is merged to a final dataset used for further steps of training or scoring (right). The coordinate system defines the surface as the *x-y* plain, and the vertical depth, perpendicular to the surface, along the red color scale *z* axis. (C) Schematic representation of the supervised machine learning classification model training. Behavior videos are manually labeled for a behavior of interest by a human observer. The XGBoost-based model is then trained with the dataset extracted by B and performance is validated for assessment of further training (blue, orange arrows) or classifier readiness (green arrow). (D) Schematic representation of automated behavior scoring. Single or multiple behavior videos go automated frame-by-frame behavior screening with trained classifiers. A frame-by-frame analysis is reported for every video together with a cohort analysis for multiple videos. Created with BioRender.

We previously reported the development of a data acquisition platform for bottom-up recordings of freely behaving mice in a dark and open field over extended periods of time, using near-infrared (NIR) light ^9^. The platform combines frame-by-frame paw-surface contact measurements and body position data for generating objective and sensitive paw luminance representations of mechanical pain and/or motor dysfunction. Nevertheless, there is also a need for the detailed quantification of specific pain-associated behavior bouts.

We introduce a tool for the automated scoring of ongoing pain-related behavior in mice ensuring easy and rapid objective quantitation. This comprises an automatic feature extraction pipeline which tracks and accurately classifies behaviors with a reliable frame-by-frame performance by analyzing animal movement in three dimensions, with a novel light-based depth feature. To evaluate this analytic platform, we focused on three ‘gold-standard’ rodent behaviors routinely used in preclinical pain research: paw flinching, licking/ biting, and grooming [8,12,16,17,33,34]. The algorithm can be trained to detect other behaviors in mice and other species in any bottom-up platform.

Recent developments in machine learning frameworks have paved direct paths to the automatic scoring of pain-related behavior from a single camera ^5^. These tools commonly use deep learning-based pose estimation tools, like DeepLabCut (DLC), to locate animal body parts in video frames, and subsequently extract the data into two-dimensional spatiotemporal features and classify the animal behavior using either supervised/unsupervised machine learning methods (e.g., support machine vectors, random forest classifiers)^10–17^.

To enhance the existing two-dimensional spatiotemporal data collection and analysis platforms, we introduce a new method for analyzing behavioral data in three dimensions by detecting changes in body part image brightness resulting from the reflection of bottom-up projected light, indicating the vertical depth of different body parts. This enables capture of three-dimensional aspects of behavior, which is more sensitive than a two-dimensional analysis, and increases the throughput of pain-related behavior analysis in a reliable, rapid and interpretable manner.

## Methods

### Animals

Adults (10 to 15-week-old) mice (male and female) were acquired from Jackson Laboratory (Bar Harbor, ME), and housed in standard clear plastic cages with 5 animals or less per cage under controlled conditions (lights on 07:00-19:00; humidity 30%-50%; temperature 22-23°C) with *ad libitum* access to food and water. All experiments were conducted between 9:00 and 17:00 in a room kept at a temperature of 21 ± 1°C. All procedures were approved by the Boston Children’s Hospital’s Institutional Animal Care and Use Committees and are consistent with NIH guidelines. For TRPV1(Cre+)-DTA experiments, wild-type C57BL6/J or TRPV1-Cre^+/+^ mice (B6.129-Trpv1tm1(cre)Bbm/J) were crossed to generate heterozygous TRPV1-Cre^+/-^ animals. The heterozygous TRPV1-Cre animals were bred with Rosa-diphtheria toxin fragment A (DTA) mice (B6.129P2-Gt(ROSA)26Sortm1(DTA)Lky/J) to produce TRPV1(Cre+)-DTA animals and their TRPV1(Cre-)-DTA littermates ^18,19^.

### Behavioral platform and video recording

The recording setup, using NIR light together with frustrated total internal reflection technology (FTIR), was described in detail by Zhang et al. ^9^, however, in this study FTIR was not used. Image brightness features were extracted using image analysis tools utilizing LED light reflected changes in pixel intensity. Briefly, an 18 × 18 × 15 cm (length x width x height) black acrylic box closed on all sides except for the bottom was placed on a 25 cm square piece of 5-mm thick glass floor. A black acrylic plate was installed under the camera, to prevent entrance of light below the glass and a 25 × 25 cm black acrylic surround frame panel with an 18 × 18 cm square cutout was positioned on top of the glass floor panel to prevent light leaking to the camera from above (Figure 1A). Two separate 850-nm NIR LED strips (SMD5050-300-IR, Huake Light Electronics Co, Ltd, Shenzhen, China) were positioned horizontally 10 cm below the glass floor to provide illumination of the animals from below and positioned such that reflections from the LEDs off the sides, top, or floor of the chamber were not visible from the camera position. Power to all LEDs was provided by a 12-V DC power supply. To record the animals in the dark an NIR camera (Basler acA2000-50gmNIR GigE, Basler AG, Ahrensburg, Germany) was positioned 30 cm beneath the glass surface. Pylon viewer software (v4.1.0.3660 Basler) was used for the video recordings. The same camera settings parameters were used for each recording. Videos were acquired at 25 Hz with 1000 × 1000 pixels dimensions and then downsized to 500 × 500 pixels for analysis using ImageJ’s scaling function, specifying to average during the downsizing. The mice were habituated in the recording device for 30 minutes before video acquisition. The floor was cleaned before the video acquisition. Recordings were performed during the animals light-cycle.

### Body part tracking

Automatic body part tracking (pose estimation) was done as described in Zhang et al. ^9^. Briefly, 50 frames per mouse were randomly chosen, each measuring 500 × 500 pixels. Nine key body parts (four paws, snout, neck, centroid, tail base and tail end) were manually labeled using DLC. These were utilized to train a DLC model for mouse pose estimation. To address model errors caused by background interferences such as urine, feces, or light reflection, the training data included videos with these potential disruptions in the field of view. The trained model was applied to automatically locate all nine body parts of interest in a video.

### Manual classification and model training

#### Dataset

We collected a data set of 24 videos from 24 mice for 3-5 minutes after capsaicin (18/24 videos) or saline (6/24 videos) subcutaneous injections, which were manually labeled by an experienced investigator for pain-related behavior bouts on a frame-by-frame level. Of the 24 videos, we used 20 for training and cross-validation testing and 4 for validation. For each of the behaviors we built a training set *(X, y)* where *X^i^* represents a set of features from frame *i*, and *y^i^* is the corresponding binary label for behavior occurrence (1; true) or behavior absence (0; false) of a specific behavior within that frame (Table 1). The occurrence or absence of a behavior was manually labeled for licking/biting, flinching, and grooming, frame-by-frame. These sets of labeled data were used to train, validate and test classification models for each of the selected behaviors. Annotations were made by a trained behavior scorer and reviewed by another trained behavior scorer, on a frame-by-frame level using a custom-built MATLAB 2021b (MathWorks, Natick, MA, USA ) script. Performance was evaluated with a set of videos that were not used for the training.

**Table 1.** Features used for training classifiers and the script names.

| Feature (inputs) | Definition | units |
| --- | --- | --- |
| **Pose features** |  |  |
| Distance (bp1, bp2)<br>*Dis_[bp1]-[bp2]* | distance between two body parts.<br>Distances normalized to the maximum distance in the distance matrix. | Pixels |
| Angle (bp1, bp2, bp3)<br>*Ang_[bp1]-[bp2]-[bp3]* | angle between three body parts | deg |
| Body part probable (bp, threshold)<br>*[bp]_inFrame(p[threshold])* | body part probability > threshold (default threshold 0.8) | Boolean |
| **Kinematic features** |  |  |
| Speed after (bp, dt)<br>*[bp]_vel[dt]* | body part speed relative to previous dt frames (default dt=2) | pix/sec |
| **Brightness features** |  |  |
| Body part's pixel intensity<br>*Pix_[bp]* | body parts pixel intensity in a frame | AU |
| Pixel ratio (bp1, bp2, area size)<br>*Log10(Pix_[bp1]/Pix_[bp2])* | absolute of log10 of ratio between mean pixel intensity of two body parts | AU |
| Body part's pixel intensity's 1<sup>st</sup> derivative before<br>*\|d/dt(Pix_[bp])\|* | Absolut of 1<sup>st</sup> derivative of the body parts pixel intensity relative to previous dt frames | AU/sec |
| Pixel ratio's 1<sup>st</sup> derivative before (Pixel ratio, dt)<br>*\|d/dt(Log10(Pix_[bp1]/Pix_[bp2]))\|* | Absolute of 1<sup>st</sup> derivative of the 'Pixel ratio' relative to previous dt frames | AU/sec |

#### Data resampling

Ongoing pain behavior bouts are commonly class-imbalanced (usually less behavior-positive frames compared to behavior-negative frames). To tackle this, for training, we re-sampled the train set by taking the average value between 50% and the real proportion percentage: (positive percent + 50%)/2 positive data instances and (negative percent + 50%)/2 negative data instances ^20^. Resampling the training data to optimized ratios of false/true target labels resulted in increased performance.

#### Feature extraction and selection

A comprehensive set of interpretable features were extracted (Figure 1B; Table S1).

#### Classifier training

Previous reports show the strength of decision tree algorithms and XGBoost tree boosting for the classification of social behavior ^13,16,21,22^. We applied XGBoost to identify a repertoire of pain-related rodent behaviors comprising licking/biting, paw flinching and grooming. To assess the classifier training performance, we used the F1-score, a metric commonly used in binary classification tasks to assess performance of a model with a score ranging between 0 (lowest) to 1 (highest) for imbalanced data. The F1-score is the harmonic mean of precision and recall, providing a balanced evaluation of both false-positives and false-negatives. Preliminary trials employing XGBoost revealed the best F1-scores while still maintaining short training and prediction times. XGBoost builds an ensemble of decision trees, which capture complex non-linear relationships from the data. It performs well with large datasets as it handles many features and is less prone to overfitting. Another advantage of XGBoost is its ability to handle missing data. Therefore, XGBoost was selected over other classifiers. A grid search with a cross-validation of 5-fold, also evaluated by F1-scores, was utilized to hyper-tune the parameters of the classification model, enabling the establishment of a common parameter set for all the selected behaviors in this study (Table 2). These parameters, however, can be further adjusted for other behaviors if needed. F1-score was measure as follows:

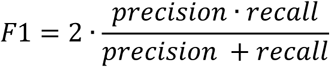

Where

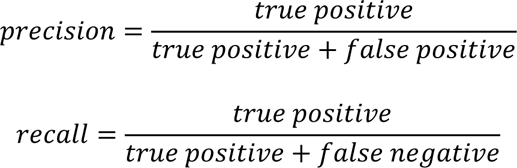

**Table 2.** Parameters used for training the classification models and prediction.

| Parameter | Values |
| --- | --- |
| **Classifier** |  |
| *n_estimators* | 1700 |
| *objective* | reg : squaredlogerror |
| *Max_depth* | 6 |
| *Learning_rate* | 0.01 |
| *Subsample* | 0.8 |
| *Colsample_bytree* | 0.2 |
| *seed* | 42 |
| *alpha* | 1 |
| *lambda* | 0.1 |
| **Postprocessing filter** |  |
| *min_bout* | Licking/biting: 4, Flinching: 5, Grooming: 1 |
| *min_after_bout* | Licking/biting: 1, Flinching: 1, Grooming: 1 |
| *max_gap* | Licking/biting: 2, Flinching: 2, Grooming: 5 |

#### Feature importance

For each behavior classifier training we applied SHAP analysis with a tree explainer algorithm, which computed the feature contribution and importance for each classifier ^23^

#### Classification threshold

For every prediction, XGBoost assigns a classification probability and makes final predictions based on a 0- to-1 threshold value, classifying instances as positive if the predicted probability exceeds the threshold, and otherwise negative. To identify the optimal probability threshold for maximizing classification performance, we implemented a k-fold cross-validation approach (k = 5) to evaluate threshold selection across folds to maximize the F1-score. For every iteration (1 of k) we shuffled data prior to splitting to reduce bias, with a consistent random state for reproducibility. Data was split into training and validation sets in an 80%/20% ratio. Thresholds were incrementally evaluated within a range of 0-to-1, in increments of 0.02, to identify the probability threshold yielding the highest F1-score for each fold. The model was fitted on the training set, and predictions on the validation set were tested across the specified range. For each threshold, we computed the F1-score, recording the threshold yielding the highest score as the optimal threshold for that fold. Following k-fold evaluations, the optimal thresholds for each fold were averaged to produce a single, cross-validated threshold representing the best overall cut-off point for classifying predictions from the model.

#### Postprocess filtering

A prevalent error in the numerous predictions we examined involved the occurrence of brief sequences (up to 5 frames) characterized by false-negatives or-false positives. To address these abbreviated error bouts, we implemented a postprocessing stage, following the primary classification. During this stage, the classifier predictions underwent a smoothing process, wherein positive instances shorter than a specified number of consecutive positive labeled frames (‘min_bout’, see Table 2), followed by a specified number of consecutive negative-labeled frames (‘min_after_bout’, see Table 2) were eliminated. Additionally, we used a gap-filling function to smooth a broken bout instance in which single frames with a bout labeled negative are switched to positive (‘max_gap’, see Table 2). These parameters can be individually optimized by users for this custom-developed classifier.

### Algorithm pipeline

ARBEL operates by receiving pose estimations data from video shots analyzed by DLC, in addition to the video file and subsequently extracts features on a frame-by-frame basis. These features encapsulate various aspects of the depicted behavior or movement. While default feature extraction parameters are available, they may also be tuned to optimize the results per organism and behavior (Table 2). ARBEL is a supervised machine learning algorithm, therefore, for training the behavior classifiers with the feature set, human scored frame-by-frame behavior targets are necessary. As a target file, ARBEL receives a comma-separated value (CSV) file type, structured with columns denoting distinct behaviors and rows corresponding to video frame numbers with each column containing frame-by-frame annotations indicating the presence or absence of the behavior. Based on these extracted features and targets, behavior classification models are generated, such as those described in the Results section. A post-analysis of threshold discrimination and a filtering process is then applied on the predicted data. The model can then be applied on a new dataset for automatic scoring.

At the end of running an experimental video analysis with ARBEL, a CSV file containing frame-by-frame automatically scored behavior labeling for each video is outputted. An additional file with total behavior duration for each behavior type is also produced. The package also contains a code for producing a video with behavior labeled frames, for convenient review and presentation.

### Data analysis computer specifications

ARBEL was implemented in Python (v3.10). It was developed in PyCharm (v2022.3.2) integrated development environment (JetBrains r.s.o, Prague, Czechia). The source code will be open and available on Github.

Machine learning analyses were performed using a consumer-grade PC with an AMD Ryzen Threadripper 3970 x 32-Core Processor, 3700 Mhz, 128 GB RAM, and NVIDIA GeForce RTX3060 GPU. All programs were run on Microsoft Windows 10 operating system.

### Reagents

Morphine Sulphate USP 2 mg/mL (Hospira, Inc., Lake Forest, IL 60045 USA) was injected intraperitoneally at 3 or 10mg/kg (10 mL/kg, in saline) or saline as control vehicle 60 minutes before the intraplantar capsaicin injection. Capsaicin (Sigma, M2028) was dissolved in ethanol to a stock solution of 0.5%. On the day of experiment, a capsaicin 0.005% working solution (diluted in saline) was prepared and injected subcutaneously into the left hindpaw in a 10uL volume.

### Statistical analysis

Statistical analyses were conducted with GraphPad Prism (v10). Unpaired t-tests (two-tailed) were employed for distributed datasets with two independent groups, while one-way ANOVAs were used for comparisons involving more than two groups to discern group effects. F1, recall and precision scores were used to compare binary datasets and Pearson correlation was used to compare non-binary datasets. Subsequently, significant main effects were further explored through post-hoc tests, specifically Tukey’s multiple comparison test. Data are expressed as mean ± SEM.

## Main

### ARBEL’s workflow from experiment to data analysis

To identify ongoing pain-related behaviors with a human-like performance we sought to employe a comprehensive approach to both capture and analyze the intricate moment-by-moment movements of mice in three dimensions. To this end, we constructed ARBEL to cater a four-step behavior-to-analysis pipeline (figure 1):

#### (1) Animal procedure and behavior recording

ARBEL is designed for a bottom-up acquisition of a control or manipulated mouse’s behavior, allowing observation of spontaneous locomotion and body part movement (Figure 1A)^9^. To capture an animal’s movements in the dark without perturbing its environment a near-infrared (NIR) LED illumination and a NIR camera was utilized (Figure 1A). Surface is not restricted to a single type of material (e.g. glass, acrylic sheets, plastic) but should be transparent, with minimal light reflectivity and camera resolution should accommodate a frame rate sufficient to detect all aspects of behavior. In this study, both glass and acrylic sheets were tested, and a camera resolution of 25Hz frame rate and 500×500 pixels were used.

#### (2) Pose-estimation and feature extraction

Following video acquisition, DeepLabCut (DLC), a marker-less pose estimation algorithm, was used to meticulously track and pinpoint the positions of key anatomical landmarks on the animal’s body and label the mice with nine body parts: snout, neck, front paws and hindpaws, centroid, tail base and tail tip of the animal (Figure 1B). This provides a high accuracy two-dimensional coordinate (*x-y* plane) for each labeled body part and their probability. Spatial (*x, y)* pixel coordinates of the DLC body part tracking data were imported into Python and processed with a custom script as part of the feature extraction process. Using the DLC output coordinates three feature types are extracted, first, spatial features that solely rely on body part *(x, y)* coordinates, including the distances and angles between them (Figure 1B, Table 1). Second, the body part coordinates from each frame are analyzed with respect to the proceeding frames to lay out the kinematic (velocity) changes of each key body part (Figure 1B, Table 1). In addition to acquiring ongoing behavior in the dark, a third, novel feature set of computer vision analysis that utilizes the NIR acquisition system to detect the animal’s body part vertical distance from the surface is extracted. Using an image analysis of the paw skin and snout, which become brighter when closer to the light source, we engineered a set of image brightness intensity-based features (Figure 1B). These features of animals’ body parts are translated into pixel intensity differences individually or compared to other body parts and go through an analysis to examine changes over time (Figure 1B, Table 1). For this study we used the left hindpaw, right hindpaw and snout, examining frame-by-frame pixel value averages for regions of interest for each frame and it’s change compared to proceeding frames (Figure 1B, green squares). To reduce the need for high computational capacity and increase running-time efficiency, light surface contact features were extracted only for three body parts in this study. However, the algorithm can be adjusted to extract spatial, kinematic, and brightness intensity features from any body part of interest.

#### (3) Training a behavior classifier

If a behavior classifier does not exist within a behavior classifier library, a new classifier can be created. This consists of manually labeling a portion of videos that include the novel behavior in a frame-by-frame positive/negative (behavior/no behavior in frame) as the training targets. ARBEL will also search for an optimal threshold using cross-validation analysis (see Methods). The threshold can be manually tuned. The extracted feature set (step 2) is used to train a classification model (‘classifier’) based on manually labeled targets. The classifier’s performance (measured with F1-score) is then tested by the user with a validation set, a set of scored videos but not used for training. A brief evaluation of the learning curve (F1-score vs. num of bout-positive frames for training) is then performed by the user. To improve the classifier: if the learning curve is inclining, more training data needs to be added to the model (Figure 1C, blue). If the learning curve reaches a plateau, model classification threshold and/or hyperparameters (Table 2) may need tuning (Figure 1C, orange) and behavior videos may need to be added. In all cases, ensuring training data includes minimal errors and representative of all variations of the behavior (pose, lighting, etc.) will increase model performance. When the classifier reaches a satisfactory performance, it is ready for automatic video analysis of unscored behavior videos (Figure 1C, green).

#### (4) Behavior analysis of a single or multiple videos

Once a behavior classifier is trained, ARBEL accepts a single or multiple videos as an input, and as an output returns a frame-by-frame behavior analysis. The video first goes through a process of pose-estimation and feature extraction. Then, behavior classification is done on a frame-by-frame basis. The algorithm returns a file containing the frame-by-frame behavioral data for each video and a screening analysis for multiple animals. A video file with the behavior classification label can be optionally produced.

##### Paw flinching classification

We tested ARBEL on its performance for classifying flinching behavior. Many instances of flinching can be missed by an observer due to their short vertical movements and fast occurrence, and it often takes more than one iteration of a behavior video examination to detect these events with high precision. To overcome this challenge, using our data, we trained a mouse paw flinching classifier (Figure 2A). To that end, we used a cohort of mice injected with capsaicin (0.005%) into their left hindpaw, which cause spontaneous flinching of the paw, and a saline injection control (Figure 2). We chose to inject capsaicin which activates TRPV1 channels expressed by a subset of nociceptors, as an acute nociceptive pain evoking chemical because of its brief (minutes) yet prominent behavioral phenotype ^24^. We tested the algorithm’s performance for flinching classification with a 5-fold cross validation F1-score learning curve. An inclining classification performance was observed between 50-to-2000 frames, in 2000 frames the learning curve reached the beginning of a stagnation point which continued in a plateau up to 8000 frames (Figure 2B). We further applied a F1-recall-precision performance test over a range of discrimination thresholds on the validation data set and found the optimal range at 0.5 (Figure 2C; threshold: 0.5, F1-score: 0.81, recall: 0.82, precision: 0.80). Next, we examined how the selected features affect the performance of the classifier. One of the advantages of gradient boosting is that after the boosted trees are constructed, the important scores for each feature for the classifier can be retrieved, allowing features to be ranked and weighted relative to each other. Feature importance analysis, using SHAP analysis, indicates the essentialness of time-dependent changes for both spatiotemporal and brightness features for the classifier’s performance ^23^. Four out of ten top weighted classifying features were image brightness-related, emphasizing the importance of these features for reaching these classification results (Figure 2d, see Table 1). Nine out of top ten features were related to the injured left hindpaw (‘hlpaw’, see Table 1). The additional tenth feature describes the velocity of the tail base in windows of 10 frames. The SHAP plot explains that higher values of tail base velocity point toward lower likelihood for flinching behavior. Indeed, flinching was scored when the animal was not locomoting.

**Figure 2.**
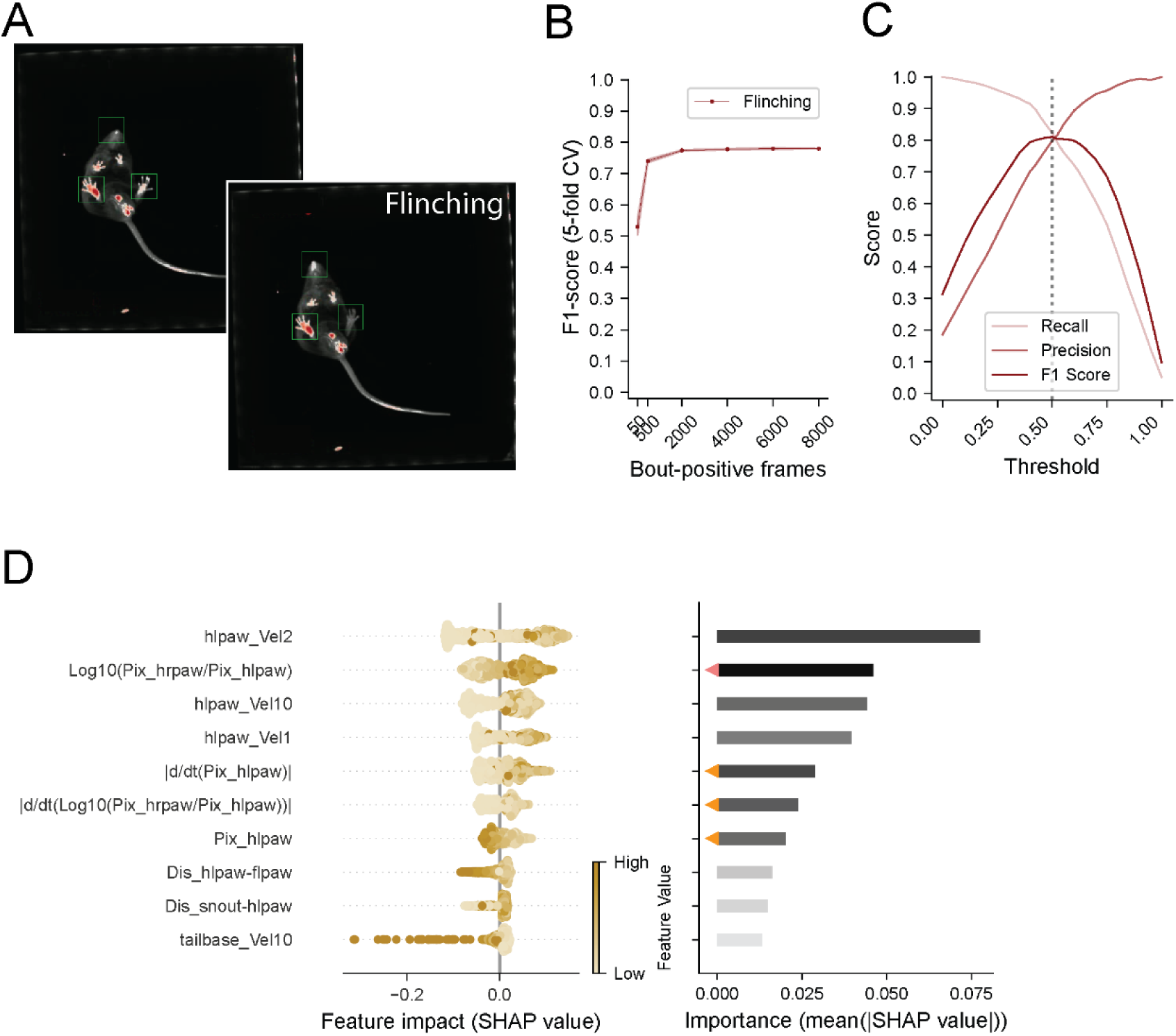
Evaluating and explaining the paw flinching behavior classifier. (A) Frames depicting paw contact before (top panel) and during flinching, overlaid with a brightness analysis (threshold > 150) (bottom panel). The green rectangles mark the regions of interest where pixel brightness ratios differentiate the right (non-injected) and left (capsaicin-injected) hindpaw. Note the difference in brightness ratio between the right and left (capsaicin injected) hindpaw. (B) F1-score-based learning curve with a 5-fold cross-validation for each data increment. (C) The model’s performance on a vaildation data set measured by F1 score (dark red), precision (red), recall (light red), and curves plotted at a 0-1 range of discrimination thresholds. The grey vertical dashed line represents the best threshold value identified during model construction. (D) Ten top important flinching classifier features for the dynamics analyzed and explained with SHAP. The impact each feature has on a bout-positive frame (positive values) or bout-negative frame (negative values) are shown on the left. Each data point represents a data point per feature out of 2000 randomly chosen and balanced features. The x position of each dot is determined by the SHAP value of that feature. Data point color is scaled based the data point value (gold scalebar) (left panel). The top ten features, from the SHAP analysis, arranged by importance weight (right panel). Orange triangles mark the brightness intensity-based features.

##### Paw licking/biting classification

Another prominent response to the induction of a localized acute nociceptive pain in rodents is repeated and ongoing licking/biting bouts of the affected paw, which are commonly monitored to quantify somatic pain after subjecting animals to acute pain (e.g., capsaicin, mustard oil) or administration of pro-inflammatory chemical irritants (e.g., complete Freund’s adjuvant, zymosan, formalin) and in models of neuropathic pain ^6–8,25–28^. We tested ARBEL for classifying ongoing licking/biting under after an intraplantar injection of capsaicin. A licking/biting bout was considered positive upon contact of the injected paw with the mouth area for two consecutive frames (Figure 3A). The learning curve evaluation showed a rapid increase between 50-to-500 bout-positive frames and this consistently graduated towards a plateau when training 2000 frames or above. (Figure 3B). Optimal discrimination threshold was set at 0.5 (Figure 3C; threshold: 0.5, F1-score: 0.91, recall: 0.92, precision: 0.90). We investigated the ten features of highest importance for classifying licking/biting behavior and found that spatial characteristics had the highest degrees of significance for this classification model (Figure 3D, see Table 1). The first ranked feature is the distance between the snout and injured left paw which indicates close contact between the paw and mouth. Interestingly, the feature importance ranking reveals that the second most important feature is the distance between the hind and front left paws. The SHAP plot further indicates that smaller values of this distance are associated with a higher likelihood of the behavior. Inspired by this finding, a closer human examination of the behavior videos confirmed that, in response to the capsaicin injection, mice bring their hind left paw toward their mouth using their front left paw during the licking/biting events. This observation highlights the physiological interpretability of ARBEL in capturing subtle nuances of behavior.

**Figure 3.**
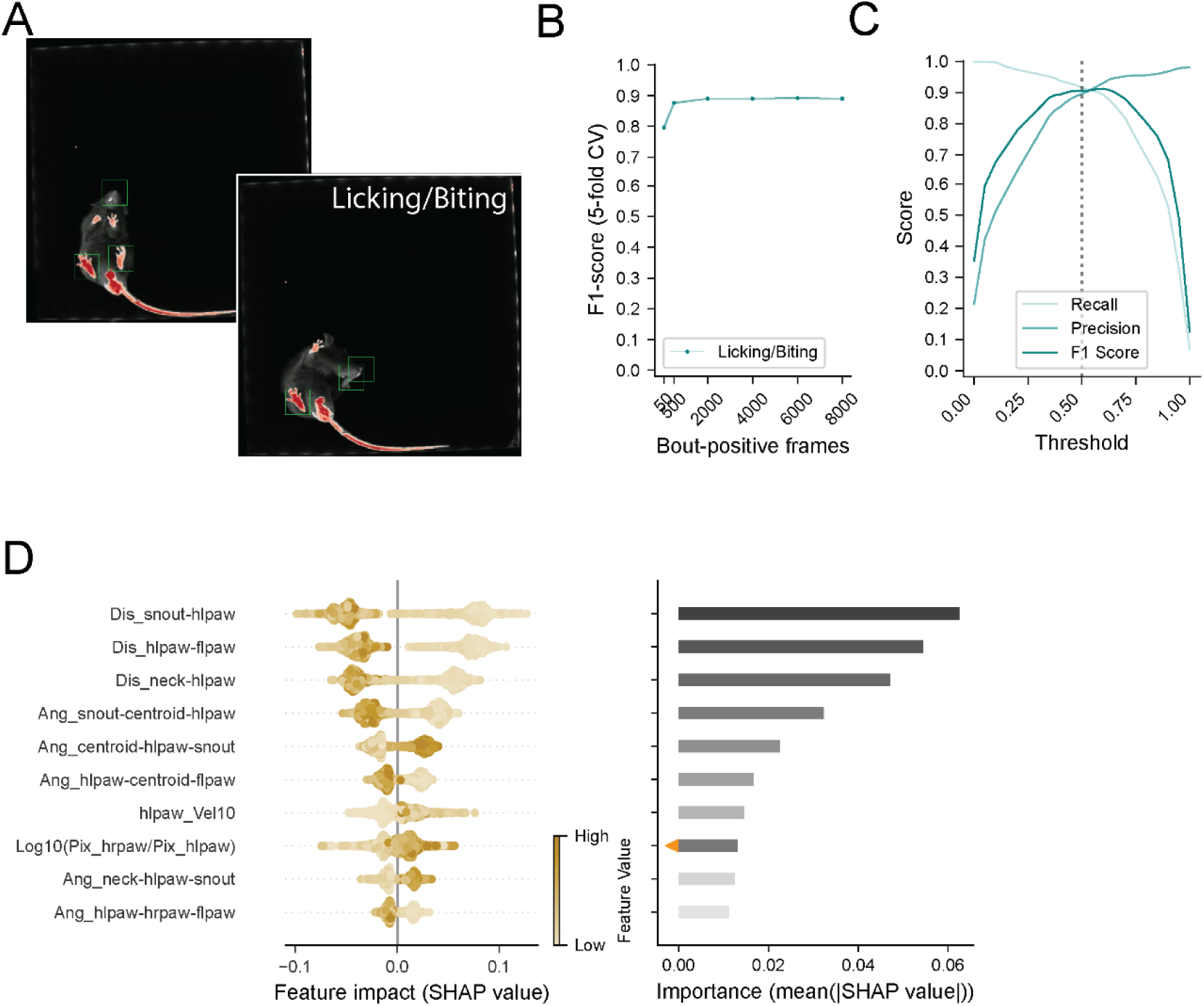
Building a classifier for paw licking/biting. (A) Frames depicting paw contact before and during grooming, with brightness analyses. The regions of interest (green rectangles) highlight the pixel brightness used to compare body-part pixel brightness interactions. (B) F1-score-based learning curve with a 5-fold cross-validation for each data increment. (C) The model’s performance on a vaildation data set measured by F1 score (dark teal), precision (teal), recall (light teal), and curves plotted at a 0-1 range of discrimination thresholds. The grey vertical dashed line represents the best threshold value identified during model construction. (D) SHAP analysis of feature importance shows the ten most impactful features for behavior classification, with their influence on bout-positive and bout-negative frames visualized (left) and their importance ranking (right). Orange triangles mark the brightness intensity-based features.

### Self-grooming classification

Rodents perform self-grooming activities throughout wakeful periods as part of normal body maintainance but this activity is increased during periods of discomfort or stress ^29–32^. Given that stress and anxiety states often coincide with acute nociceptive and inflammatory pain, changes in grooming behavior have been proposed as a potential indicator of persistent pain ^2,30,33^. Grooming can also change in response to analgesics ^34,35^. Mice self-groom various parts of their bodies, including their backs, sides, genitals, limbs and tails by licking or nibbling, or their faces using their forepaws to rub their cheeks and eyes (Figure 4A). While humans can easily recognize and categorize all these different bouts as grooming behavior, it encompasses a wide range of actions involving several different body parts. Hence, in comparison to the flinching or licking/biting behaviors, grooming demands a more mathematically complex and robust classifier to accurately capture and differentiate all the behavior nuances. We tested our algortithm for detecting grooming on the sames set of data used for training the flinching and licking/biting classifiers. Learning curve, as in previous behaviors, started its plateau at 2000 frames(Figure 4B). The threshold for the grooming classification was also found to be within optimal ranges at 0.4 (Figure 4C; threshold: 0.4, F1-score: 0.82, recall: 0.93, precision: 0.72). Interestingly, some of the top weighting features of importance involved the DLC probability of the body part visibility withing a frame (‘_inFrame_p0.8’, see Table 1)(Figure 4D). During some behaviors, some of the body parts are hidden from the camera’s view and will therefore have low probability of the pose estimator’s identification. For example, the front paws may be masked by the head during face grooming, and the snout by the body during back grooming. We found that above 0.8 is a sufficient criterion threshold indicating for whether the body part is clearly visible to the human eye.

**Figure 4.**
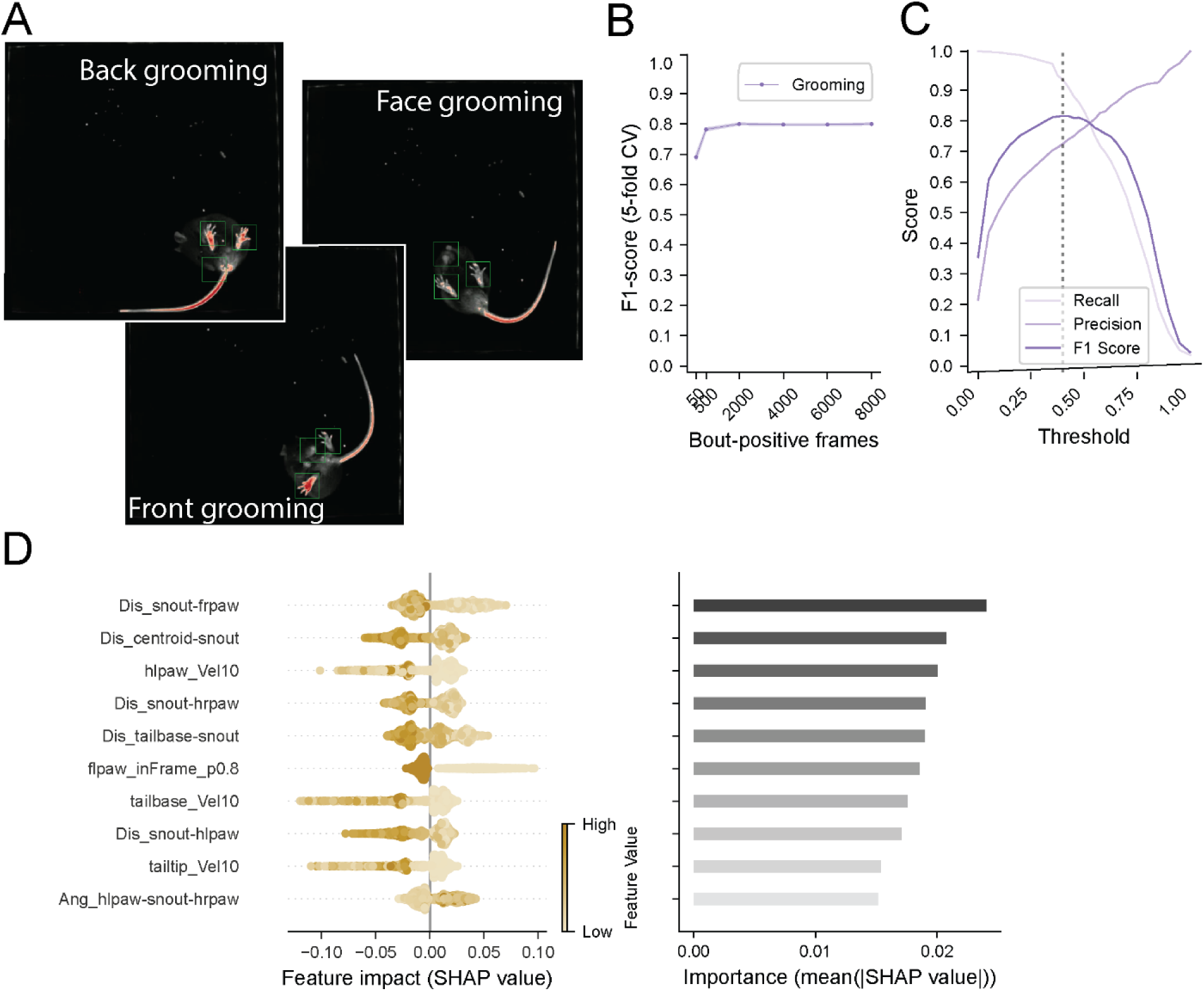
Grooming behavior classifier exploration. (A) Representative frames before and during various grooming behaviors. The green rectangles depict the region of interest in which pixel brightness is averaged for quantifying paw contact with the surface. Note the difference in brightness ratio between the right and left (capsaicin injected) hindpaw. (B) F1-score-based learning curve with a 5-fold cross-validation for each data increment. (C) The model’s performance on a vaildation data set measured by F1 score (dark purple), precision (purple), recall (light purple), and curves plotted at a 0-1 range of discrimination thresholds. The grey vertical dashed line represents the best threshold value identified during model construction. (D) SHAP analysis shows the ten most impactful features for behavior classification, with their influence on bout-positive and bout-negative frames visualized (left) and importance ranking (right).

### Classification model testing

To evaluate performance we tested the classifiers on a continuous three-minute test video recording (4500 frames), of a mouse’s behavioral response to injection of capsaicin into its left hindpaw, while comparing it to with an experienced manual scorer’s labeling. The model performed well in identifiying flinching and licking/biting and behaviors, with an F1-score of 0.82 and 0.90 respectively, on a frame-to-frame resolution (Figure 5A). It’s grooming behavior prediction also performed with a high human versus model F1-score of 0.86 (Figure 5A). We further examined the model’s performance for total licking/biting, flinching and grooming prediction durations over time in 5 second time bins and found that at this resolution the prediction was well aligned with the human expert labeling, reaching 0.89, 0.99 and 0.97 respectively with Pearson correlation (*R*) (Figure 5B).

**Figure 5.**
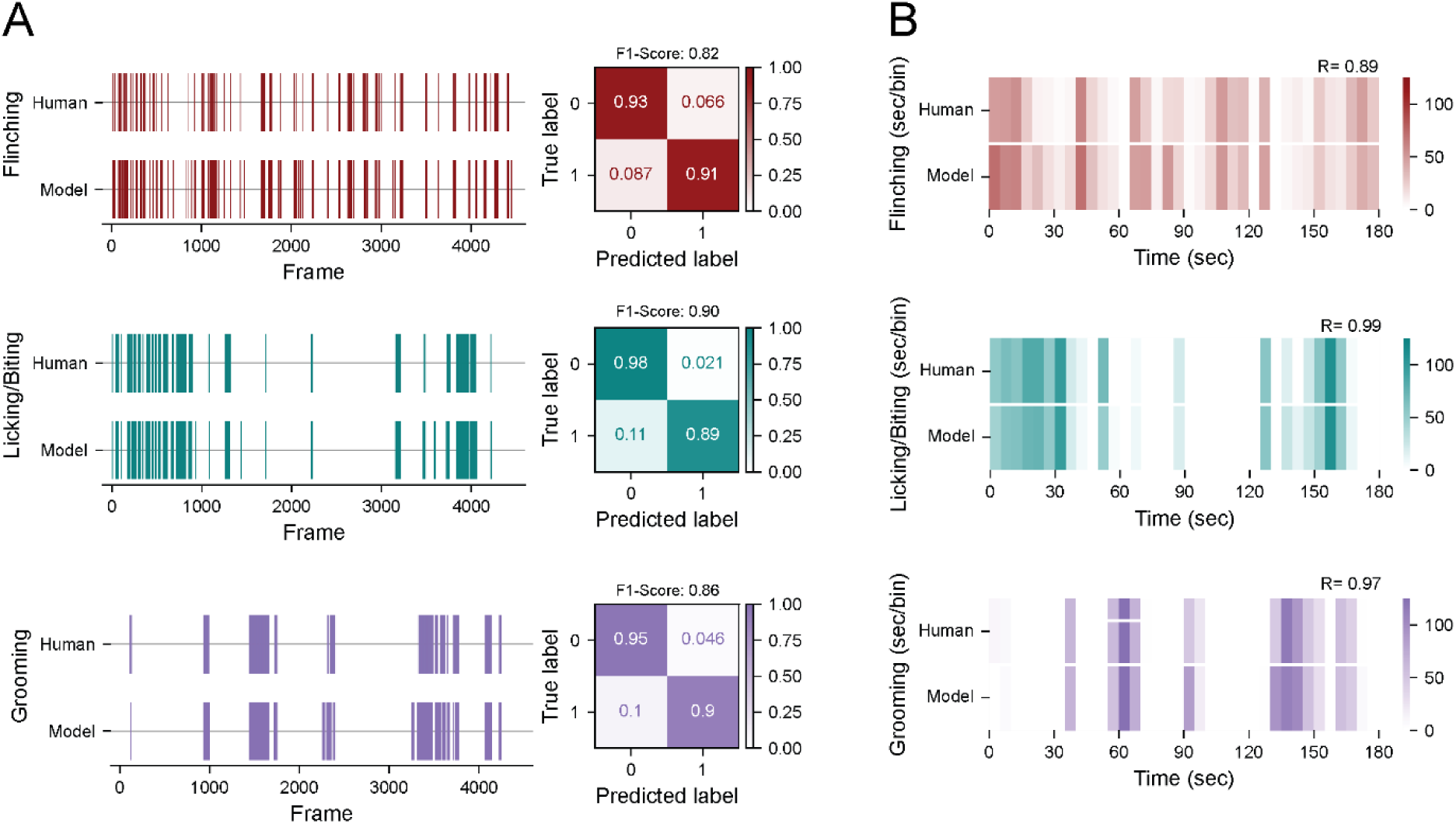
Testing pain-behavior and grooming classifier performances. (A) Examples of three-minute frame-by-frame (25Hz, 4500 frames) ethograms of human annotation versus the classification model’s predicted bouts of flinching (red), licking/biting (teal), and grooming (purple) and their corresponding F1-scores and confusion matrices. (B) Ethograms displayed in five-second time bins with Pearson’s correlation test score (*R*) on the top right.

### Automatic behavior screening of analgesic drug efficacy and genetic mutation

To evaluate the algorithm’s utility for screening differences in behavior in large datasets aimed at detecting drug efficacy or genetic mutations, we administered subcutaneous capsaicin to five cohorts of mice and recorded their behavior over a 3-minute period. Mice were pretreated intraperitoneally (i.p.) with vehicle, 3 mg/kg, or 10 mg/kg morphine one hour prior to capsaicin injection. Additionally, behavioral responses were screened in TRPV1(Cre+)-DTA mice, in which TRPV1 neurons were ablated, and their TRPV1(Cre-)-DTA littermates. Changes in flinching behavior revealed the clear dose-dependent effect of morphine. Mice pretreated with 3 mg/kg morphine did not exhibit significant reductions in flinching compared to vehicle-treated controls. However, the 10 mg/kg dose significantly reduced flinching duration relative to both the vehicle group and the 3 mg/kg group. Similarly, TRPV1(Cre+)-DTA mice displayed significantly less flinching compared to their TRPV1(Cre)-DTA littermates (Figure 6A). Paw licking/biting behavior was significantly reduced in mice pretreated with 3 mg/kg morphine compared to vehicle-treated controls, with an even greater reduction observed in the 10 mg/kg group. This dose-dependent analgesic effect highlights the sensitivity of the algorithm in capturing subtle behavioral changes. TRPV1(Cre+)-DTA mice also exhibited markedly less licking/biting compared to TRPV1(Cre-)-DTA mice, validating the role of TRPV1 neurons in mediating this nociceptive behavior (Figure 6B). Grooming behavior was effectively abolished in both the 3 mg/kg and 10 mg/kg morphine groups, with no significant differences between the two doses. These findings align with previous reports of morphine-induced suppression of grooming behavior, demonstrating the robustness of the algorithm in detecting established drug effects. Unlike flinching and licking/biting, grooming behavior did not significantly differ between TRPV1(Cre+)-DTA and TRPV1(Cre-)-DTA mice, suggesting it is less influenced by TRPV1-mediated nociception (Figure 6C). These findings underscore the robustness of the algorithm in unraveling nuanced, dose-dependent behavioral responses and its efficiency in segregating genetic models by phenotype. By enabling the simultaneous analysis of multiple behavioral endpoints over continuous observation periods, the algorithm provides a powerful tool for the preclinical screening of analgesics and validating genetic models of nociception.

**Figure 6.**
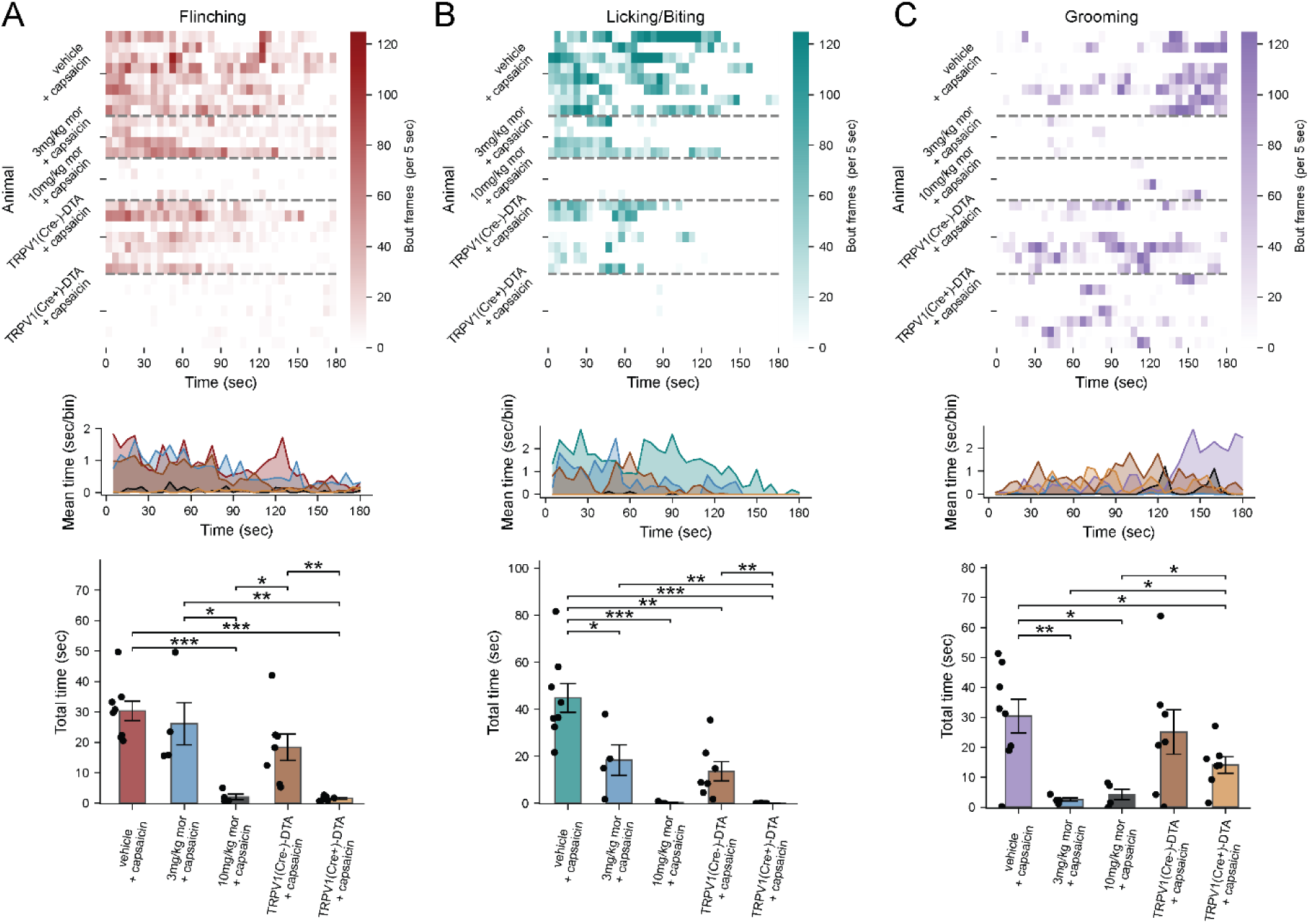
Automated behavior screening for analgesic efficacy and genetic models of nociception after injection of capsaicin. The algorithm’s detection of dose-dependent effects of morphine on flinching, licking/biting, and grooming behaviors following subcutaneous capsaicin administration in mice. (A) Top, heat map representing flinching time in 5 sec bins. Groups of screened mice included wildtype mice, pre-treated (i.p.) vehicle (n=8), 3mg/kg morphine (n=4) or 10mg/kg morphine (n=4), naïve TRPV1-(Cre+)-DTA (n=7) and TRPV1-(Cre-)-DTA (n=7) mice. Middle, Summary of total flinching time in the heatmap per animal group. Bottom, quantification and statistical analysis of total flinching time for each group of mice. Cohorts are separated by grey dashed lines. (B,C) Same as (A) for licking/biting and grooming respectively. Data represented as mean ± SEM. Statistical significance determined by student’s T-test (* p<0.05, ** p<0.01, *** p<0.001). Black circles, single animals.

## Discussion

This study introduces an innovative analytic approach to identify and quantify ongoing pain-related behaviors in mice, employing an interpretable supervised machine learning algorithm integrated with a bottom-up open field animal recording platform ^9^. The combination of the existing hardware and the new software achieves performance levels matching human scoring but in a fraction of the time required, offering reliable and efficient means of detecting ongoing pain-related behaviors.

Humans identify and classify behaviors based on subtle cues and patterns that may be difficult to quantify mathematically. For example, a flinch is recognized due to its abrupt nature or a paw lick due to the animal’s body posture and its repetitive motion. However, defining these behaviors mathematically involves consideration of variables such as distances between body parts, movement speed, trajectory, duration, and context, among others. Each of these variables, or ‘features’, can interact in complex ways, making the creation of a clear, objective, and comprehensive mathematical definition a challenging, and in many cases unfeasible task. Here we implement a machine learning tool to train classifiers on labeled human-annotated behaviors to learn those intricate patterns and nuances that clearly characterize each type of behavior from the data itself. The classifier, once trained, effectively embodies a definition of the behaviors it was trained to recognize, although in a form that is derived from data rather than from explicit rules. Furthermore, using SHAP analysis we show that physiologically explainable features of behaviors can be detected using ARBEL’s features set.

We also introduce a novel approach for collecting hindpaw skin and snout brightness data derived from video analyses of NIR reflection signals, in addition to a DLC-based spatiotemporal dynamics feature set. The enhanced performance of this behavior screening platform with the open-source software provided, opens new possibilities for efficient and accurate behavioral screening in mice with a temporal element^9^. This approach enables an acceleration of the pace of experimentation and reduces variability which has the potential to increase the scope, efficiency and reproducibility of analgesic drug discovery initiatives. By facilitating a comprehensive assessment over extended periods, the platform is a valuable tool for detecting behavioral changes in inflammatory or chronic pain models and for detecting the effects of therapeutic interventions. Furthermore, it is also beneficial for screening genetic models contributing to detecting those genes engaged in driving pain-related behaviors.

In this study we only highlighted flinching, together with licking/biting and grooming behavior classifiers. Flinching and licking/biting are directly associated with ongoing pain behavior and are considered gold-standard indications for pain in rodents, with flinching considered a defensive response, and licking/biting a coping response^6,7,36^. Although grooming is not commonly considered an indicator of pain, it serves as an important measure of stress, anxiety, and discomfort, clear side effects of pain or of adverse effects of analgesics like morphine ^34,37,38^. By unraveling the intricate interplay between drug dosage and changes in behavior one can gain insight into desired actions or side effects that manifest in a behavior-specific manner. Although we chose here to showcase ARBEL’s utility for scoring three pain-related behaviors the classifiers can be trained for other behaviors (Figure 1C)

Stimulus evoked-pain methods, such as Von Frey monofilaments or Hargreaves heat radiance tests at single time points are widely used to quantify pain in preclinical studies, since the results are immediate unlike ongoing behavioral measures which necessitate prolonged observation^4^. Devices that standardizes and automates pain testing by providing computer-controlled aiming, stimulation, and precise response measurement with thermal, mechanical and optogenetic stimuli have been recently introduced^39,40^. Automation of stimulus-evoked pain assessments is essential to increase the pace and precision of drug discovery. However, ongoing inflammatory or neuropathic pain in a naturalistic movement environment, is more clinically relevant for drug efficacy profiling. Our algorithm’s ability to differentiate and quantify pian-related behaviors may increase the clinical relevance of preclinical pain phenotypes and the evaluation of new therapeutic strategies.

ARBEL is not restricted to interpreting behavior only with the behavioral platform used in this study, or only in mice since it can be customized for any video data set that can be interpreted with DLC. Although we show the advantages of our approach for generating information on ongoing pain initiated in a hindpaw, this is not obligatory for training a custom behavioral classifier. The software can be tuned and trained for any set of pose estimation outputs, from diverse organisms. Our findings show that for mice the hindpaw and snout brightness ratio is a valuable feature supporting classification of the three behaviors measured in this study, but users can adapt and tune its application to any labeled body part. The brightness signals which indicate distance from the surface improves detection and quantitation of certain behaviors, however the utilization of the brightness features may not be required for all classification performance.

Users may seek to acquire insight into social interactions between subjects. While the current iteration of the algorithm’s classifiers is tailored to handle a single animal, a multiple animal scenario can readily be addressed by running multi-animal pose-estimation, as offered by DLC and other pose estimation algorithms, and train classifiers for the designated multi-animal behaviors using the same feature extraction process detailed here (Figure 2; see Methods) ^12,41^.

Although the primary focus of this study was on automatic pain-related behavior scoring, the approach has the potential to be adapted to other behaviors associated with and require detailed ongoing observation like stress, anxiety, locomotor disorders, and neurological degeneration and developmental disorders. Another prospect will be to integrate the bottom-up behavioral platform and ARBEL’s automatic behavior scoring with *in-vivo* neuronal activity monitoring techniques such as electrophysiology, electroencephalogram (EEG), fiber photometry, or single-cell mini-scope recordings with fluorescent molecular genetic tools in freely behaving animals. This would help elucidate specific neuron-behavior couplings and help provide a comprehensive understanding of cell-to-network levels of neuronal activity during observed behaviors. This could also enable automatic real-time closed-loop experiments, allowing excitatory/inhibitory manipulation of specific neural circuits (e.g., channelrhodopsin2 stimulation) in response to behavioral states of interest.

We optimized our code to operate within the constraints of standard computer specifications but used a GPU recommended for deep learning pose estimation. A fundamental aim was to ensure that the algorithm is both simple and flexible, allowing for easy coding adjustments in a Python software environment. While numerous behavior analysis programs have been published, this algorithm introduces a novel approach, specifically designed to leverage changes in body part brightness to extract useful three-dimensional features for the precise evaluation of animal pain-associated behavior. ARBEL’s goal is to complement and enhance existing tools by providing a robust and innovative way of precisely and automatically detecting and assessing an animal’s pain-related behavior. Its application can advance analgesic treatment development, as well as any preclinical research associated with behavioral outputs, by reducing labor, time, and animal usage, and increasing the objective and quantitative determination of the behavioral consequences of changes in the peripheral and central nervous system.

## Acknowledgements

NIH: R35NS105076 and R01AT011447(CJW) and Animal Behavior Core 5P50HD105351, DARPA: HR0011-19-2-0022 (CJW).

